# Pioneering the Formation of 2-Carboxylic Anthraquinone: CRISPR/Cas9-Mediated Functional Validation of *Octaketide Synthase* and *Polyketide Reductase* Genes in *Aloe vera*

**DOI:** 10.64898/2026.01.12.698956

**Authors:** Alka Jangra, Siddharth Tiwari, Vinod Chhokar

## Abstract

*Aloe vera* is an authentic medical plant abundant in aromatic polyketides, including the crucial hexaketides aloenin, aloesin, and barbaloin used in pharmaceuticals yet the enzymatic basis of their biosynthesis remained incompletely understood. While it has been suggested that octaketide synthases (OKS) initiates anthraquinones biosynthesis, heterologous expression of OKS alone consistently produces shunt polyketide products, and the mechanism underlying this derailment was uncertain. To comprehend the mechanism of anthraquinone biosynthesis, we combined biochemical constitution, structural characterization and CRISPR/Cas9-mediated editing of key genes in *Aloe vera*. It was for the first time demonstrated that the inclusion of a PKR (polyketide reductase) redirected the reactive intermediate toward formation of 2-carboxy anthraquinone (C_16_H_12_0_5_). The identity of reaction product was confirmed by spectroscopic analysis which additionally rendered it clear from compounds previously misannotated in the literature. Alongside, CRISPR/Cas9-based genome editing of *OKS* and *PKR* genes resulted in significant reduction (upto 2.54 fold) in aloin content in edited lines compared to the non-edited control aloe line. Together these findings endorsed the presence of tailoring enzyme ketoreductase for the efficient and appropriate formation of anthraquinones and establish a mechanistic framework for polyketide biosynthesis in aloe plants that sustain as an indigenous herb for mankind.

**Highlights:** This study provides direct biochemical and genetic evidence that a tailoring enzyme is required to prevent derailment of polyketide intermediates and enable correct anthraquinone formation in plants.

## Introduction

Plants have served as an imperative source of medication for many years, and almost 80% of the world’s population still depends upon natural means of remedies for their healthcare. *Aloe vera* enjoys a tremendous global demand and acceptance due to its pervasive medicinal, nutraceutical, and many other uses (Rajeswari et al., 2012). *Aloe vera* Linné (*Aloe barbadensis* Miller) is medicinally most potent and, consequently, more prominent (Babu and Noor, 2020; Al-Dulaimi, 2022). Until now, the literature reports various biochemical, transcriptomics, and metabolomics analyses of the aloe plant and its application in curing various health disorders such as asthma, diabetes, skin problems, microbial infection, etc. (Choudhri et al., 2018; Jangra et al., 2022, 2023a). These medicinal properties of the plant are attributed to more than 75 active secondary metabolites it retains, including various efficacious polyketides and secondary metabolites like anthraquinones, alkylbenzene, polysaccharides, lectin, lignin, saponins, etc. (Sharon and Suvarna 2017; Abe 2020). Aloin, an anthrone C-glycoside, is a modified anthraquinone with an extra sugar molecule attached *via* C-C linkage and is an abundant anthraquinone in *Aloe vera* plants (Patel and Patel, 2013; Lewis et al., 2022). Aloin A and aloin B (two isoforms of aloin) are reported to selectively inhibit the proteolytic and deubiquitinating activity of PLpro (Papain-like) protease of SARS-CoV-2 (Lewis et al., 2022). However, aloin administrations have been reported to increase severities and incidences of persistently active inflammation in the large intestine, and its toxic effects could be mild to severe depending upon the dosage and route of intake (Boudreau et al., 2017). Under the National Toxicology Program conducted by the National Cancer Institute in 1998, *Aloe vera* was nominated for tumor promotion and carcinogenicity based on its high oral and tropical consumption by men, women, and children (Jangra et al., 2022).

The co-action of chalcone synthase (CHS), a type III polyketide synthase (PKS) with octaketide synthase (OKS) activity, and polyketide reductase (PKR) portrays a crucial function in the synthesis of diverse polyketides and several antimicrobial compounds in both leguminous and non-leguminous plant species (Bomati et al., 2005). OKS is hypothesized to catalyze the biosynthesis of anthraquinones in plants by facilitating the sequential condensations of eight molecules of malonyl-CoA and the products often spontaneously fold into complex aromatic/cyclic compounds (Abe et al., 2005a; Guo et al., 2021; Møller et al., 2016). PKRs are crucial supplementary enzymes in numerous plant polyketide biosynthesis pathways comprising the aldo-ketoreductase (AKR4) family in plants (Morita et al., 2007; Krishnamurthy et al., 2022). It is evident that the accessory PKRs are essential for stabilizing the linear intermediates and prohibiting them from undefined cyclization during their production in the lactone-producing pathways initiated with PKS (Zhu *et al*., 2022). Previous studies revealed the formation of unphysiologically folded polyketides (SEK4 and SEK4b) from heterologously expressed (*E. coli*) *OKS* of *Aloe arborescens.* SEK4/SEK4b and their metabolites are considered shunt products of the OKS pathway and are not naturally accumulated in plant tissues. Their formation depends upon the presence/absence of downstream enzymes/partners (Abe et al., 2005a; Møller et al., 2016; Kang et al., 2020). Since the role of OKS and PKR in polyketide biosynthesis remains to be elucidated, it is compelling to hypothesize their role in the biosynthesis of anthrones and anthraquinones in aloe. The present study inflated the in-depth understanding of *OKS* and *PKR* genes involved in the aloin/anthraquinones biosynthesis pathway in *Aloe vera*. *De novo* transcriptome sequencing of *Aloe vera* provided unigene sequences encoding OKS and KR, likely to be involved in anthraquinone biosynthesis in aloe (Choudhri et al., 2018). Building upon the transcriptome dataset, we have trailed the apprehension of *OKS* and *PKR* genes by heterologous gene expression in *E. coli* and CRISPR/Cas9 (Clustered Regularly Interspaced Short Palindromic Repeat and associated protein) system to substantiate their role in the biosynthesis of aloin.

## Material and method

### Plant material, vectors and bacterial strains

*Aloe vera* (*Aloe barbadensis* Miller) plants were raised in the nursery of Guru Jambheshwar University of Science and Technology, Hisar (India). The DH5α strain of *E. coli* was utilized for the cloning process, while the BL21 strain (*E. coli*) was employed for expressing heterologous genes. To investigate the sub-cellular localization of the candidate gene in *Nicotiana benthamiana* leaves, the GV3101 strain of *Agrobacterium tumefaciens*, carrying the binary vector pCAMBIA1302, was used. All cultures were grown in Luria-Bertani Broth (LB) medium. Enzymes used in this study were sourced from New England Biolabs (NEB, Ipswich, Massachusetts, USA). The RevertAid First Strand cDNA Synthesis Kit (ThermoFisher Scientific), was used for cDNA synthesis. Various antibiotics, including gentamycin (100 µg/ml), ampicillin (100µg/ml), rifampicin (50 µg/ml) (Sigma-Aldrich, St. Louis, Missouri, USA) and kanamycin (50 µg/ml) (Himedia, Thane, Maharashtra, India) were employed. All chemicals utilized were of analytical grade and chromatographically pure.

### *In-silico*, physicochemical, and phylogenetic analysis of *OKS* gene Identification and physicochemical analysis of *Aloe vera* derived *OKS* gene

The *OKS* (*Locus_3851_Transcript_1/9*) was retrieved from the previously reported leaf transcriptome dataset of *Aloe vera* available at Sequence Read Archive (NCBI) *via* accession number SRR5167034 (https://www.ncbi.nlm.nih.gov/sra). The conserved domain analysis of identified *OKS* was performed by the NCBI conserved domain database (CDD) and Simple Modular Architecture Research Tool (SMART) domain analysis database. The physiochemical analysis was carried out on Protparam (https://web.expasy.org/protparam/). Subcellular localization of identified protein was predicted by Cello 2.0 (http://cello.life.nctu.edu.tw/) (Yu *et al*., 2004). The multiple sequence alignment (MSA) of protein sequences was performed with Clustal X 2.0 (http://www.clustal.org/clustal2/).

### Phylogenetic analysis of OKS protein

To attain the maximum understanding of the evolutionary relationship among *Aloe vera* and different dicots and monocots species, the phylogenetic tree was constructed. For OKS, protein sequence of various monocot and dicot plant species **(Table S1)** were retrieved from Ensembl plant genome database (https://plants.ensembl.org/index.html) based on similarity search. The phylogenetic tree was constructed using the pairwise deletion method in MEGA X software (Tamura et al., 2021) with the neighbor-joining algorithm and default parameters, including 1000 bootstrap replicates, Poisson correction distance, and pairwise deletion. The circular tree was further refined using iTOL (https://itol.embl.de/upload.cgi).

### Functional characterization of *OKS* gene by recombinant protein expression in *E. coli* Identification and cloning of *OKS* gene

Total RNA was extracted from the healthy young leaves of *Aloe vera* plants grown in the nursery of Guru Jambheshwar University of Science and Technology, located in Hisar, Haryana, India. The isolation of RNA was performed using the LiCl precipitation method. Subsequently, the isolated RNA was converted into complementary DNA (cDNA) following the instructions provided in the cDNA synthesis kit. For heterologous expression of *OKS* gene, gene specific primers having customary restriction site at their ends were designed **(Table S2)**. These primers were designed on the basis of *Locus_3851_Transcript_1/9* coding sequence (CDS). The primers were engineered with *Nde*I at the 5’ end, *BamH*I at the 3’ end for OKS sequences. With the gene-specific primers, the candidate gene was amplified from cDNA of *Aloe vera* using Hotstart Phire DNA II polymerase (Thermo Fisher Scientific, USA). The purified *OKS* gene was cloned into the pJET 1.2 cloning vector. After confirmation, the recombinant plasmid was digested with the corresponding restriction enzymes and the eluted gene fragment was then cloned into linearized pET28a bacterial expression vector.

### Gene expression and protein purification

A single colony from the positive transformants of OKS gene was cultivated in LB containing 50 µg/ml kanamycin at 37°C, 150 rpm until OD_600_ reached 0.6 and then subjected to induction using 0.5 mM IPTG (isopropylthio-β-galactoside). The induction of OKS protein was carried out at 28°C and 150 rpm for 8 hours. Both induced and un-induced, were harvested at 7000 rpm, 4^°^C for 15 min, and processed immediately or stored at -30^°^C till further use. The pET-28a vector carries an N-terminal His tag, and an optional C-terminal His tag sequence that was utilized to purify protein *via* Ni-NTA affinity purification. The purified protein was run on SDS (sodium dodecyl sulphate) polyacrylamide gel (12%) and visualized through coomassie brilliant blue staining. The purified proteins were further concentrated by dialysis. The concentration of purified and concentrated protein was determined by Micro BCA Protein Assay Kit (GENETIX).

### Enzymatic activity of OKS enzyme with the starter substrate

The functional characterization of OKS was performed through an enzymatic reaction with malonyl-CoA. The enzyme assay of OKS was performed as per the reaction described by Abe et al., (2007) with some modifications. The reaction mixture included 216 μM of malonyl-CoA and 10 μg of recombinant OKS enzyme, in a final volume of 500 μL of 100 mM potassium phosphate buffer with a pH of 6.0. The incubation was carried out for 1 hour at 30 °C. The resulting products were extracted with 1000 μL of ethyl acetate and filtered through 0.22 µm membranes (HiMedia, India) into amber color HPLC vials. The sample was sonicated for 15 mins in an ultrasonic cleaner (Aczet Pvt. Ltd., India) and analyzed using Agilent Poroshell 120 EC-C18 ( 3.0x100 mm, 2.7 μm particle size, 120 Å pore size) HPLC column (Agilent Technologies, USA) at a flow rate of 0.8 mL/min. The HPLC gradient elution was performed using a mixture of H_2_O and MeOH, both containing 0.1% TFA. The elution gradient protocol was as follows: 0-5 mins, 30% MeOH; 5-17 mins, 30-60% MeOH; 17-25 mins, 60% MeOH; 25-27 mins, 60-70% MeOH; 27-35 mins, 70% MeOH; 35-40 mins, 70-100% MeOH.

### Cumulative activity of OKS and PKR enzymes

The catalytic activity of PKR enzyme was conducted using the established standard assay conditions as reported in our previous report (Jangra et al., 2023a). To study the cumulative action of OKS and PKR, a reaction mixture was prepared as described by Oguro et al., (2004) with some modifications. The reaction contained 216 μM of malonyl-CoA, 1 mM NADPH, and 5 μg of recombinant OKS enzyme, along with 60 μg of recombinant PKR enzyme (Jangra et al., 2023b), in a final volume of 500 μL of 100 mM potassium phosphate buffer with a pH of 6.0. The incubation was carried out for duration of 2h at 30°C, and the reaction was stopped by adding 50 μL of 20% HCl. The resulting products were extracted with 1 mL of ethyl acetate and analyzed using reverse metabolomics. The ethyl acetate extract of the reaction was evaporated using SpeedVac system (Thermo Fisher Scientific, Waltham, MA, USA) and analyzed *via* ESI-MS in the positive ion mode. Further analysis was carried out using Fourier-transform infrared spectroscopy (FTIR) and Nuclear magnetic resonance (^1^H NMR). FTIR and MS spectra of Chrysophanol analytical standard (Sigma-Aldrich, Merck, USA) was also procured for structural correlation and comparison.

### Functional characterization of *OKS* and *PKR* genes in anthraquinone biosynthesis pathway using CRISPR/Cas9 based genome editing

#### Selection of gRNA target sequence and construct preparation

A target-specific single guide RNA (gRNA) was generated individually for *OKS* and *PKR* genes using the full-length CDS sequence *Aloe vera* using the CRISPR-P v2.0 tool (http://crispr.hzau.edu.cn/CRISPR2/) (LeiY, 2014) and a 20-bp target sequence with a protospacer adjacent motif (PAM) at the 3’ end was selected for targeting the *OKS* and *PKR* genes **(Table S3)**. A pair of target gRNA was selected for each gene that showed maximum efficiency score and cloning linkers were synthesized for each gene from the Eurofins genomics (Bengaluru, India). The synthesized oligonucleotides were annealed and phosphorylated to generate an oligo duplex, and ligated into the *Bsa*I-digested pRGEB31 vector following the instruction provided in the cloning guide for pRGE vectors (Addgene, MA, USA). The ligated modified vector was transformed into competent *E. coli* (DH5α) cells using heat shock method and the recombinant plasmid was isolated from the transformed *E. coli* clones using the GeneJET Plasmid Miniprep Kit (Thermo Fischer Scientific, USA) and confirmed with restriction digestion followed by sequencing. The positive CRISPR constructs were further transformed into the AGL1 strain of *Agrobacterium tumefaciens* and designated as CRISPR-OKS and CRISPR-PKR.

#### Transformation, selection, and regeneration of *Aloe vera*

The CRISPR constructs, CRISPR-OKS and CRISPR-PKR, were transformed into the pre-cultured shoots of *Aloe vera via Agrobacterium* following a standard protocol optimized by adjusting various parameters, as reported in our previous study (Jangra et al., 2023a).

#### Screening of transformants and mutation study

Total RNA was isolated from the leaves of putative edited and control (non-edited) lines of *Aloe vera* and reverse transcribed into cDNA following the user instructions. The putative transgenic plants were further checked for the presence of *Cas9* by using *Cas9* specific primers **(Table S4).** To check the target mutation in the gene sequence of the *OKS* and *PKR* genes in putative edited aloe lines, a set of primers were designed for each gene to amplify the desired DNA fragment. The amplified fragment was isolated from the gel using the GeneJET Gel Extraction Kit (Thermo Fischer Scientific, USA) and directly sequenced using a 3730xl DNA analyzer (Applied Biosystems, USA). The deduced nucleotide sequence data was aligned with the gene sequence of the *OKS* and *PKR* genes using Multalin software (http://multalin.toulouse.inra.fr/multalin/) to determine the mutation pattern and the effect of the mutation was predicted by using Expasy Translate (https://web.expasy.org/translate/).

#### Aloin estimation in edited and non-edited control *Aloe vera* lines

The whole leaves were harvested from the edited aloe lines of *OKS* and *PKR* genes as well as from the non-edited control set of plants for estimation of their aloin content. The biological triplicates were considered, and three different plants from the same gene were used for the sampling. The aloin content was estimated in whole leaf extract (WLE) of the edited and non-edited aloe lines using a previously reported procedure with some modifications (Kumar *et al*., 2016). The methanol extract of the sample was dried at 45°C for 24h in a hot air oven and dissolved in extraction buffer (1mg/ml) containing 35% acetonitrile (ACN, HPLC grade) and 65% H_2_O (HPLC grade). The samples were filtered through 0.22 µm membranes (HiMedia, India) into amber colour HPLC vials and sonicated for 15 mins in an ultrasonic cleaner. The aloin content was evaluated using ZORBAX Eclipse Plus C18 (4.6x250 mm, 5 μm particle size, 70 Å pore size) HPLC column (Agilent Technologies, USA). A mobile phase of 35% ACN (HPLC grade) and 65% H_2_O (HPLC grade) in an isocratic condition with a flow rate of 0.6 mL/min and 5µl injection volume was used. Aloin values were measured in micrograms per mg of fresh weight (µg/mg, FW), and quantification was carried out at 210 nm with reference to the aloin standard (Sigma-Aldrich, USA). Chromatography data integration was performed using Mass Lynx software.

### Sub-cellular localization of *OKS* gene by transient expression in *Nicotiana benthamiana*

cDNA synthesized from the total RNA of *Aloe vera* was used to amplify the *OKS* gene using gene-specific primers. The primers used for the expression of *GFP* at the N and C terminal of the gene were designed and amplified using the NEBuilder® HiFi DNA Assembly tool **(Table S5)** and Master Mix (NEB). The fragment amplified with N-terminal primers was digested using the *Sfu*I restriction enzyme, while the fragment amplified with C-terminal primers was digested with the *Nco*I restriction enzyme and cloned into the backbone of vector pCAMBIA1302. The positive clones were introduced into the GV3101 strain of *Agrobacterium tumefaciens*. The infiltration of *Nicotiana benthamiana* leaves with the pCAMBIA-OKS(N) and pCAMBIA-OKS(C) constructs (Fig. 3), and pCAMBIA empty vector was carried out according to the methodology of Thakur et al., (2021) with some modifications. In brief, the recombinant cells of *Agrobacterium* were cultured and suspended in the MMA medium comprising 0.1 M MES buffer (pH: 5.2), 0.3 M MgCl_2_, and 0.2 M acetosyringone (1-(4-hydroxy-3, 5-dimethoxy phenyl) ethan-1-one) corresponding to 0.8 OD at 600 nm. The mixture was incubated at 28°C for 6-8 h with gentle shaking and consequently availed for infiltration.

The wild-type *Nicotiana benthamiana* L. (tobacco) plants were cultivated in growth chambers (Conviron, Winnipeg, Manitoba, Canada) maintained at 24±2°C under 16/h photoperiod with a light intensity of 250 μmol m^-2^s^-1^ provided by cool-white fluorescent tubes (standard culture conditions). The leaves from 4-6 week old plants were infiltered with *Agrobacterium* MMA suspension using a 2 ml syringe (without any needle). The *Agrobacterium* cells comprising non-targeted GFP (pCAMBIA1302 empty vector) were used as a control vector to curtail the non-specific localization. The plants were incubated in the dark after infiltration for 24h under moist conditions and thereafter transferred to the growth chamber under standard conditions. The GFP fluorescence was detected under the confocal microscope.

### Statistical analysis

All the experiments in the present study were performed by using two to three biological replicates and three technical replicates. Student’s paired t-test and Dunnett’s one-way variance (ANOVA) were performed according to the experiments that are displayed as mean ± SD. The mean values for each treatment were compared (*P ≤ 0.05; **P ≤ 0.001; ***P ≤ 0.0001) to assess significance levels. GraphPad Prism 5 software was used for the data analysis.

## Results

### Transcriptome mining identifies candidate *OKS* and *PKR* genes involved in anthraquinone biosynthesis in *Aloe vera*

The nucleotide and deduced amino acid sequence of the *OKS* and *PKR* gene were submitted to the NCBI database (https://www.ncbi.nlm.nih.gov/) *via* GenBank accession number BankIt2714563 Locus_3851_ Transcript_1/9 OR146495 and BankIt2620826 AvAKR, espectively (Jangra *et al*., 2023b). The conserved domain analysis of OKS protein showed the presence of a catalytic active site (Cys^174^, His^316^, and Asp^349^), malonyl-CoA binding site, along with a product binding site, collectively representing the conserved domain of chalcone synthase superfamily **(Fig. S1)**. Furthermore, MSA of OKS deduced amino acid sequence also showed the presence of conserved catalytic triad, in accordance with the previously reported sequences of the respective genes from other plant species **(Fig. S2)**. The physicochemical analysis of OKS protein revealed the presence of a 1212 bp long coding region encoding 403 residues and a molecular weight of 44.28 kDa, where each residue holds an average weight of 109.87 Da. The isoelectric point of the sequence was determined to be 5.85, signifying its theoretical pH at which it carries no net charge. Additionally, the sequence displays a minimal probability of approximately 0.62% for expression in inclusion bodies. Moreover, the sub-cellular localization of *OKS* was predicted to be cytoplasmic, which was later confirmed with the Green Fluorescent Protein (GFP) tag in tobacco leaves.

### Phylogenetic analysis indicates close relatedness of aloe-based OKS to monocot PKS

The phylogenetic analysis in the present investigation was intended to investigate the evolutionary associations of aloe-based candidate genes with other crop species. According to the phylogram, the OKS enzyme derived from *Aloe vera* is closely related to the members of the monocot group, such as *Aloe arborescens, Asparagus officinalis*, *Zea mays*, and *Avena sativa*. These species share a common evolutionary lineage with the aloe OKS enzyme. In contrast, the other major clades of dicot species appear to be more distantly related to the OKS clade **(Fig. 1)**. This observation revealed the taxonomical similarity with monocot crops, while other species exhibited distant genetic relations with putative OKS from *Aloe vera*.

**Fig. 1.**
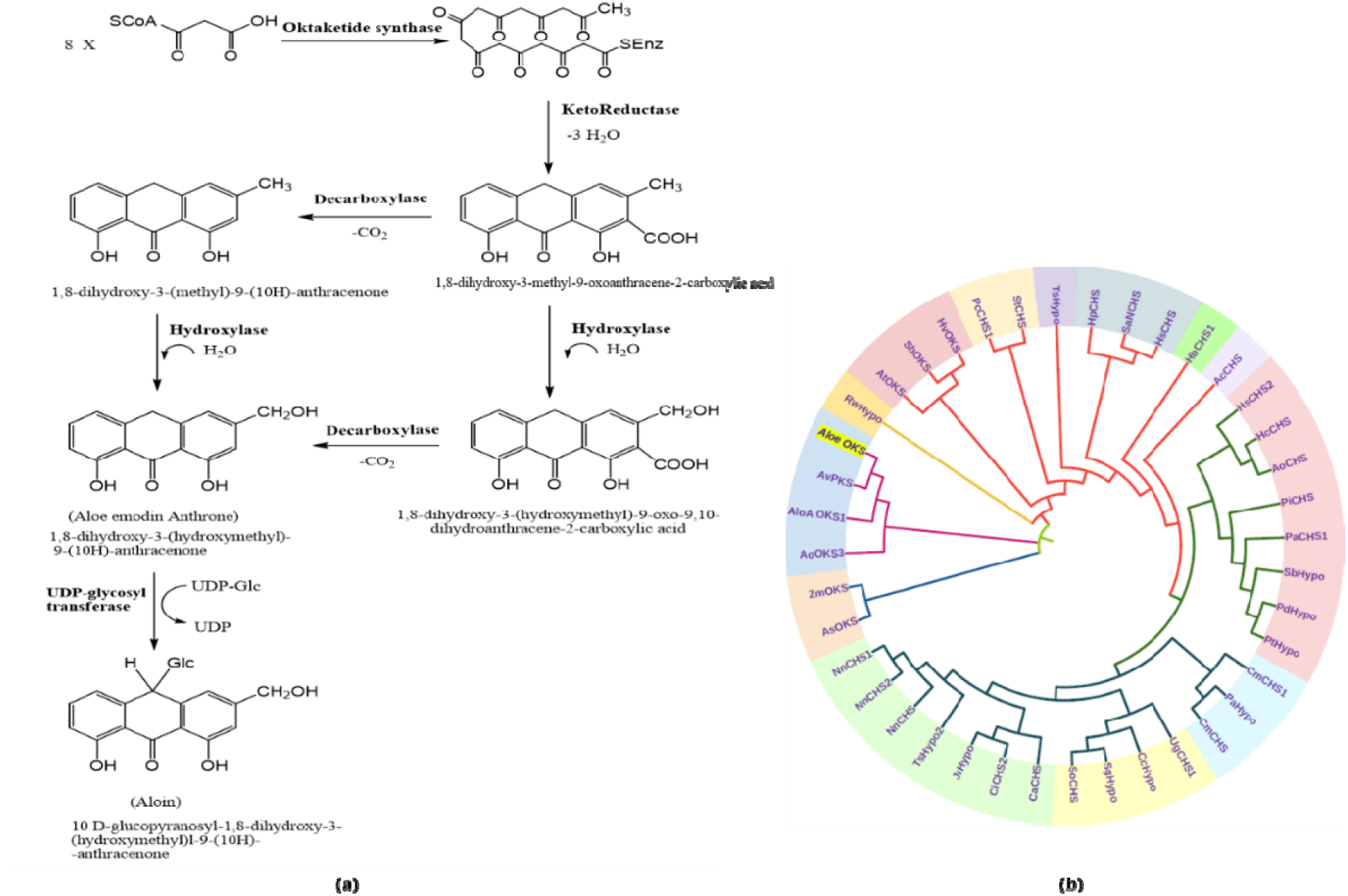
Identification of *OKS* and *PKR* genes involved in aloin biosynthesis in *Aloe vera;* (a) Proposed pathway illustrating the hypothesized role of OKS and PKR in directing anthraquinone formation; (b) Phylogenetic analysis showing clustering of *Aloe vera* OKS with other dicot and monocot species, indicating evolutionary conservation. The bootstrap values were calculated with 1000 replications

### Characterization of OKS enzyme: Recombinant OKS alone generate shunt products

The CDS of the *OKS* gene was amplified and successfully cloned into pET28a (+) and transformed into BL21 expression cells of *E.coli*. The SDS-PAGE analysis confirmed a strong expression of OKS protein, mostly in the soluble form. A considerable band of about 49 kDa was observed on SDS-PAGE gel for OKS protein. The protein expressed by the recombinant cells was a fusion protein containing a linked His-tag of approximately 5 kDa. Further, western blot analysis validated the positive binding of recombinant protein with His-tag antibody (Fig. 2a). The recombinant OKS enzyme utilized the starter substrate malonyl-CoA and resulted in the synthesis of SEK4/SEK4b **(Fig. 2c)**, the longest polyketide ever formed by a structurally simple homodimeric type III PKS. Retention time: SEK4 (12.7 min) and SEK4b (14.7 min).

**Fig. 2.**
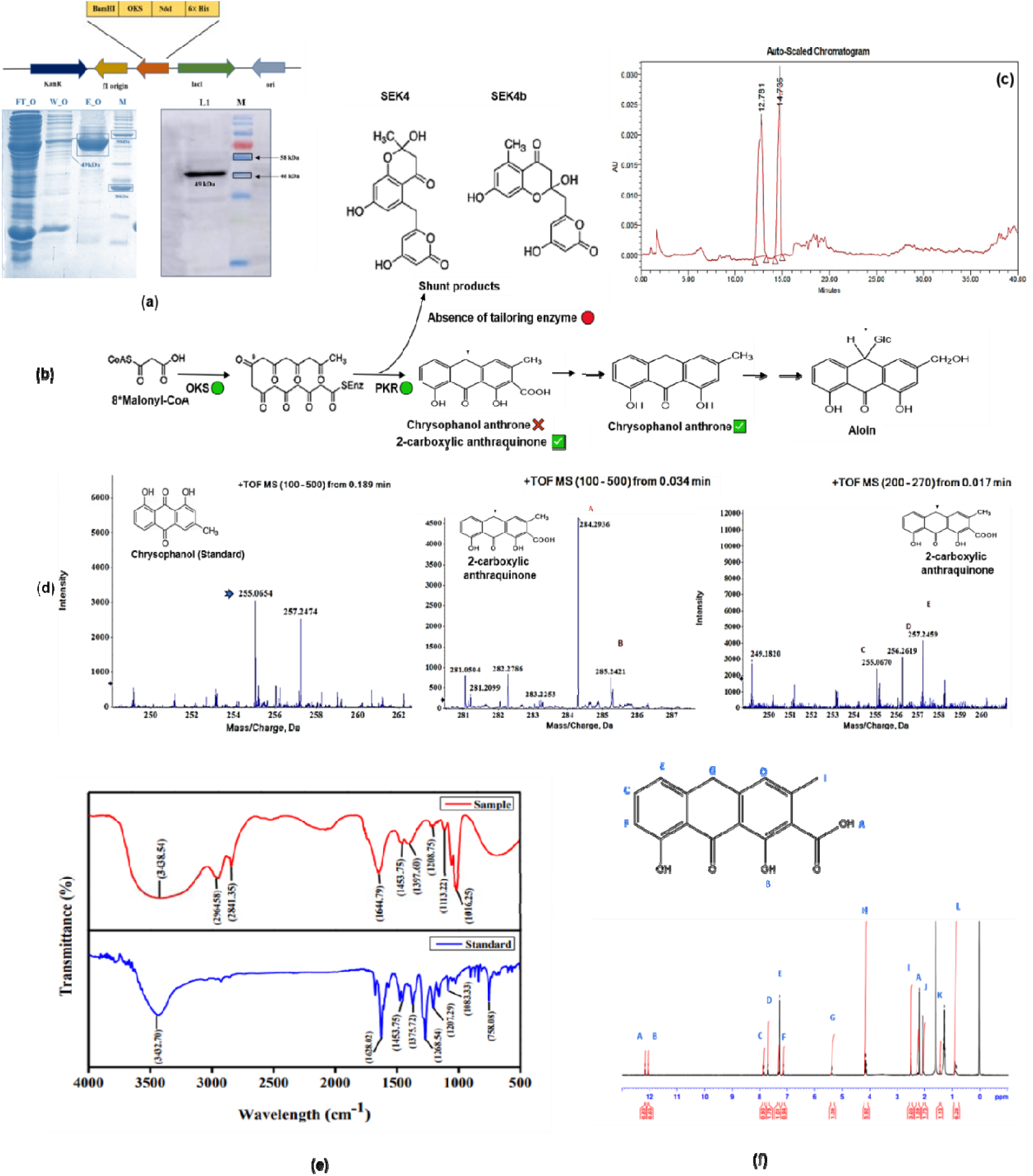
Fig. Biochemical reconstitution of anthraquinone biosynthesis reveals cumulative action of OKS and PKR; **(a)** SDS–PAGE and western blot analyses showing soluble expression and purification of recombinant octaketide synthase (OKS) and polyketide reductase (PKR) proteins used for enzymatic assays; **(b)** Proposed model for aloin biosynthesis in *Aloe vera* and distinguishing it from compounds previously described as chrysophanol anthrone **(c)** HPLC chromatogram of the in vitro reaction catalyzed by OKS alone using malonyl-CoA as substrate, showing accumulation of the shunt products SEK4 and SEK4b, indicative of aberrant cyclization in the absence of a tailoring enzyme; **(d)** ESI-MS spectrum of the OKS–PKR reaction product showing molecular ion peaks corresponding to 2-carboxy anthraquinone ([M]C and [M+H]C) as compare to chrysophanol; **(e)** FTIR spectra of the OKS–PKR reaction product showed diagnostic bands associated with hydroxyl and carboxylic acid functionalities, consistent with the formation of 2-carboxy anthraquinone and distinct from chrysophanol; and **(f)** Structural confirmation of the OKS–PKR reaction product by ¹H NMR spectroscopy

### Cumulative action of OKS and PKR enables formation of the native anthraquinone scaffold

To comprehend the mechanism of the aloin biosynthesis, the present study investigated the enzymatic conversion of the malonyl-CoA by recombinant aloe based OKS and PKR together. It was for the first time demonstrated that the malonyl-CoA was converted to 1,8-dihydroxy-3-methyl-9-oxoanthracene-2-carboxylic acid (2-carboxylic anthraquinone) by the co-action of the OKS/PKR enzyme system. This investigation initially focused on examining the mass spectrometric characteristics of the product formed, considering its chemical structure and the similarities it shares with the previously reported compounds and their metabolites. The analysis of the product *via* ESI-MS (SCIEX TripleTOF 5600 and 5600+ /SCIEX) in the positive ion mode is shown in **Fig. 2d**. The molecular cation (2-carboxylic anthraquinone, C_16_H_12_O_5_) was detected by ESI-MS in [M]^+^ and [M + H]^+^ forms (m/z 284.2936 and 285.2421). The parent compound also showed fragment ions of m/z 255.0670 ([M + H − CO]^+^) and 257.2459 ([M + H – C_2_H_4_]^+^). The distinctive fragmentation pattern exhibited by the parent compound’s product ions was utilized to aid in its characterization. For instance, the ion exhibiting m/z 255.0670 ([M + H − CO]^+^) is identical as chrysophanol as per the ESI-MS spectra of chrysophanol standard.

Further analysis with FTIR showed significant O–H stretching indicated by a broader peak at 3412 cm^−1^ as compared to the chrysophanol standard suggesting the presence of a characteristic carboxylic group in the product formed. The assay extract also marked significant peaks at 2950.94 and 2843.54 cm^−1^, along with shoulder peaks at 2981.21 and 2923.88 cm^−1^ (aliphatic C-H). The FTIR spectra of the sample and chrysophanol standard exhibit similar infrared features as follows: well-pronounced peaks around 1670 to 1760 cm^−1^ (aromatic C=C, H-bonded C=O), sharp peaks around 1452 cm^−1^(CH_2_), and peaks in 1150 to 1160 cm^−1^ region (aliphatic CH2, OH or C-O stretch of various groups). The spectra of the product differ from those of chrysophanol in several sharp, small peaks and shoulders exhibited in 2800 to 3500 cm^−1^region owing to the presence of H-bonded carboxylic group and anonymous aliphatic chain comprising aliphatic C-H region (Inbar *et al*., 1989; Singh *et al*., 2013), thereby suggesting the formation of 2-carboxylic anthraquinone as a cumulative product of OKS and PKR **(Fig. 2e)**. The IR spectra of the samples were recorded on Perkin Elmer Spectrum, BX II (version 10.6.02) spectrophotometer in KBr tablets.

The ^1^H NMR spectra was recorded at 400MHz on Bruker, Advance III spectrophotometer in CDCl_3_ revealed signals at δ_H_ 12.161 and 12.054 ppm, indicating the presence of strong acidic groups, such as carboxylic acid (COOH) or a strong acidic proton, five ortho-coupled aromatic protons (δ_H_ 7.85, 7.70, 7.28, 7.13 ppm) with coupling frequency (J) 9.6Hz and 10.8Hz for doublets at 7.85 ppm and 7.70, respectively, one proton associated with an unsaturated carbon (δ_H_ 5.36 ppm, J-4.4Hz), six protons (δ_H_ 4.14 ppm, J-6.8Hz) associated with functional group or a saturated carbon (CH_2_ or CH_3_) group, chemical shifts at δ_H_ 1.2 (J-7.2Hz), 2.0, 2.1, and 2.4 ppm suggesting the presence of eight protons associated with aliphatic or saturated carbon atoms, and around six protons (δ_H_ 0.84 ppm, J-6Hz) associated with a terminal methyl group (Fig. 2f).

### CRISPR/Cas9-mediated mutagenesis of OKS and PKR genes revealed their role in aloin biosynthesis

The CRISPR-OKS and CRISPR-PKR constructs were successfully prepared using the binary vector pRGEB31 and the verified plasmids were effectively transformed into Agrobacterium cells (AGL1). The individual recombinant Agrobacterium cells carrying the CRISPR-OKS and CRISPR-PKR constructs were used to transform aloe shoots. After undergoing a strict selection process, independent shoots were obtained for each construct. The putative transformed shoots were transferred onto rooting media, then acclimatized in a potted Soilrite mix. The plantlets were successfully acclimatized in growth chambers before being moved to the greenhouse for further growth and development.

### Frame shift mutations result in premature truncation of OKS and PKR proteins

To confirm the transformation of aloe plants, the DNA of putative transgenic plants was amplified using *Cas9*- gene-specific primers. The transformed transgenic plantlets exhibited the desired band size of 190 bp **(Fig. 3b)**, which was consistent with the positive control.

**Fig. 3.**
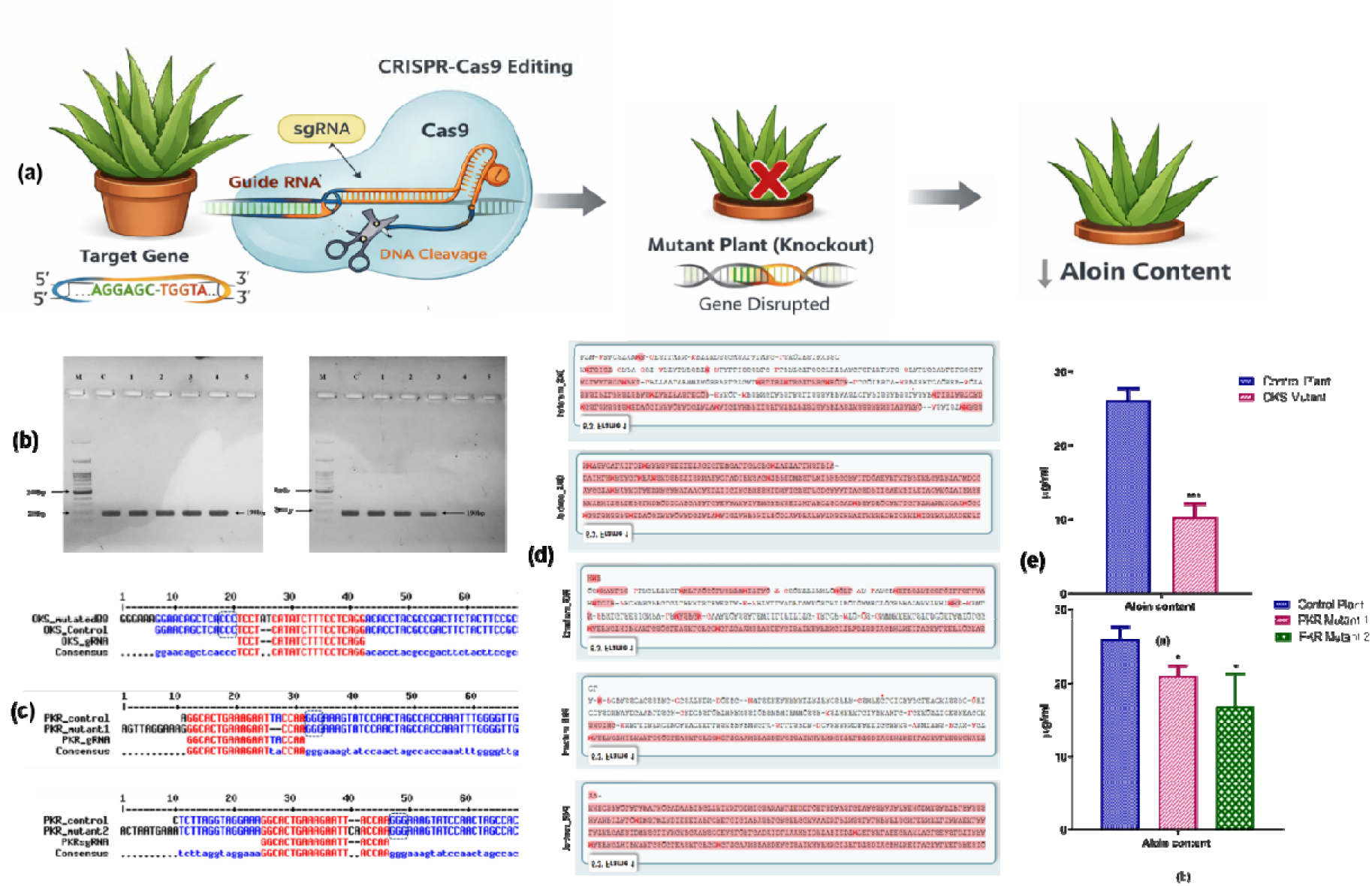
CRISPR/Cas9–mediated functional validation of *OKS* and *PKR* genes in *Aloe vera*; **(a)** Schematic illustration of CRISPR–Cas9–based genome editing in *Aloe vera*, showing sgRNA-guided cleavage of the target gene resulting in gene disruption and reduced aloin accumulation; **(b)** CRISPR–Cas9 editing of the OKS/PKR gene showing the sgRNA target region and PAM sequence, with Cas9-induced mutations occurring within the sgRNA recognition site; **(c)** Representative sequence alignment and translated protein view of wild-type and edited alleles showing frame shift mutations within the coding region, resulting in truncated proteins; and **(d)** Quantitative analysis of aloin content in non-edited and CRISPR-edited *Aloe vera* lines, demonstrating a significant reduction in aloin accumulation upon disruption of OKS or PKR

Most of the putative transgenic lines amplified appropriate bands, indicating a positive PCR result. Following the PCR screening for positive lines, further screening was conducted through sequencing to confirm the editing of the *OKS* and *PKR* genes. The sequencing results revealed the editing of the *OKS* gene sequence only in 1 transformed aloe line, while 2 lines accorded the editing of the *PKR* gene sequence. The results showed an insertion of 2 subsequent bases AT and CA upstream of the protospacer adjacent motif (PAM) sequence individually, in one transformed line of OKS and one transformed line of PKR (mutant 2), respectively. While deletion of 2 two consecutive bases, TA was accorded in one transformed line of PKR (mutant 1) **(Fig. 3c)**. *In-silico* analysis further revealed that these double base insertions and deletion in different lines caused frame shift mutations in the target region, leading to the disruption of protein synthesis and the possible knock-down function of *OKS* and *PKR* genes **(Fig. 3d)**.

### Loss of function in *OKS* and *PKR* genes leads to a significant reduction in aloin content in edited lines

Aloin estimation was carried out in WLE of both edited and non-edited control aloe plants. There was only one edited line of CRISPR-OKS and two edited lines of CRISPR-PKR constructs. The results showed that the WLE of all three edited lines possessed lower aloin levels as compared to the non-edited control plants **(Fig. 3e)**. The control lines contained 26.09 µg/mg of aloin, while the edited line of CRISPR-OKS encompassed 10.27 µg/mg (OKS mutant) of aloin in WLE. The aloin content in WLE of two PKR edited lines was accorded to be 21.16 µg/mg (PKR mutant 1) and 16.98 µg/mg (PKR mutant 2), respectively. On average, the CRISPR-OKS edited lines had 2.54 fold lower aloin content than the non-edited control aloe line, while the CRISPR-PKR edited lines contained 1.23 and 1.53 fold lower aloin content in mutant 1 and mutant 2, respectively.

### OKS localizes predominantly to the cytoplasm with peripheral association to the plasma membrane in tobacco cells

The putative *OKS* gene were successfully amplified, cloned in pCAMBIA 1302 plant expression vector with a GFP tag **(Fig. 4)**, and infiltered in tobacco leaves. The transiently expressed fusion protein was visualized using confocal microscopy, which revealed a distinct subcellular localization of fluorescence in cells expressing the GFP in tobacco leaves. The fluorescence defining the expression of untargeted GFP accumulated in the cytoplasm, while the fluorescence representing the fused GFP with OKS (pCAMBIA-OKS at C-terminal as well as N-terminal) was found to be localized in the cytoplasm and lining the plasma membrane, suggesting the cytoplasmic localization of the candidate gene.

**Fig. 4.**
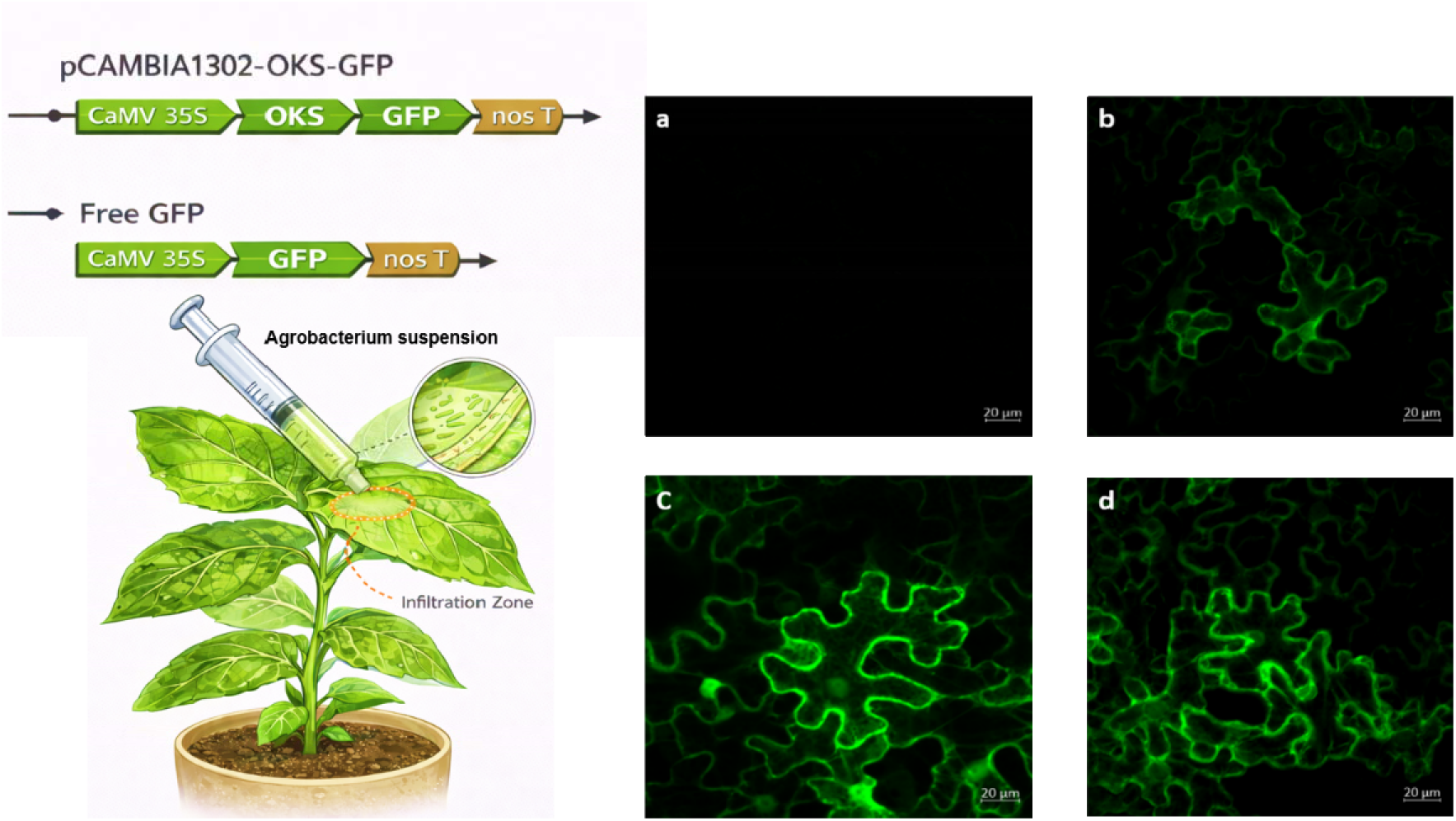
Schematic illustration of Agrobacterium-mediated transient expression and subcellular localization of OKS in tobacco leaves. General vector map of pCAMBIA-OKS (N-terminal) construct used for the transient expression of *OKS* gene and *Nicotiana benthamiana* L. (tobacco) leaves. Transient expression of GFP fused-OKS in **(a)** control plant leaf, **(b)** expression of control untargeted GFP **(c)** expression of PKR-GFP construct (C-terminal), and **(d)** expression of OKS-GFP construct (C-terminal)

## Discussion

The present investigation provides direct biochemical and genetic evidences that aloin/anthraquinone biosynthesis in *Aloe vera* require cumulative activity of *OKS* and downstream tailoring enzyme, *PKR*. The study employed heterologous expression of *OKS* in *E. coli* and CRISPR/Cas9-mediated editing of *OKS* and *PKR* in *Aloe vera* to comprehend this mechanism. The *OKS* (Locus_3851_Transcript_1/9) and *PKR* (unigene_74658) sequences were retrieved from the previously reported leaf transcriptome dataset of *Aloe vera*. The physicochemical analysis and domain analysis of OKS protein representing the conserved domain of the CHS superfamily. This is in line with previous studies, where structural and mutational analyses of plant type III PKS enzymes revealed that the conserved catalytic triad (Cys, His, Asp) is fundamental to their decarboxylation-condensation reactions during polyketide chain elongation. The presence of conserved active-site residues in MSA emphasizes the significance of these residues in facilitating catalytic reactions and highlights their functions in the synthesis of complex molecules in biological systems (Pandith et al., 2020; Abe et al., 2005a; Jez et al., 2000).

While OKS has long been proposed as the initial enzyme in polyketide intermediate formation, the inability of OKS alone to produce the native anthraquinone scaffold has remained unresolved. The formation of SEK4 and SEK4b by the OKS enzyme is consistent in many previous investigation, which reported the formation of these aromatic octaketides by *Aloe arborescens*-based PKS (Abe et al., 2005a). The formation of these byproducts by the type III PKSs was described as "non-catalytic" cyclisations of the nascent polyketide. These cyclisations occur in the absence of a specific ketoreductase and are different from the typical type III PKSs based catalytic cyclisation reactions. The minimum PKS created the major ring of SEK4, which corresponds to the C7–C12 section of the octaketide, without the involvement of a ketoreductase (Fu et al., 1994a,b). Several plant-specific type III *PKSs*, including *PcOKS* (Guo et al., 2021), *DlHKS* (heptaketide synthase) from *Drosophyllum lusitanicum* (Jindaprasert et al., 2008), *PiHKS* (hexaketide synthase) from *Piriformospora indica* (Springob et al., 2007), *HsPKS1* from *Huperzia serrata* (Wanibuchi et al., 2007), and *WtPKS1* from *Wachendorfia thyrsiflora* (Brand et al., 2006) were expressed in *E. coli* but did not generate the expected products. Transient or hyperexpression of *Aloe arborescens* OKS (Aa*OKS*) in *Nicotiana benthamiana* also formed SEK4 and SEK4b. However, when co-expressed along with Streptomyces sp. *R1128 cyclase* genes *in vivo* production of different downstream compounds such as flavokermesic acid anthrone was detected. Similarly, the transgenic lines of *Nicotiana benthamiana expressing* PzPKS produced naphthalene derivatives, in the absence of Plumbago-specific downstream enzymes (Jadhav et al., 2014; Møller et al., 2016). In *Senna tora,* CHS-L9 produced anthranoid scaffolds and early intermediates *in-vitro*, but not final oxidized anthraquinones (e.g. emodin), also indicating the absence of additional downstream enzymes (oxidation, decarboxylation, etc.) (Kang et al., 2020). Cumulatively, these findings implies that it is possible to convert the non-reduced octaketide of first step into C_14_-C_35_ aromatic aglycon compound of interest subjected to the presence of tailoring enzyme (Møller et al., 2016).

Our findings demonstrate that PKR is essential for preventing derailment of reactive polyketide intermediates and enabling formation of 2-carboxy anthraquinone, establishing a mechanistic basis for pathway fidelity in this medicinal plant. It was proved for the first time that the malonyl-CoA was converted to 2-carboxylic anthraquinone by the co-action of the OKS/PKR enzyme system. The molecular cation (2-carboxylic anthraquinone, C_16_H_12_O_s_) was detected by ESI-MS in [M]+ and [M + H]+ forms (Prateeksha et al., 2019). The structure formulated with FTIR and 1H NMR spectral data also propounds the existence of an anthraquinone molecule possessing a carboxyl, hydroxyl, and terminal methyl group (Inbar et al., 1989; Singh et al., 2013). These spectral data and the formularized structure indeed suggest the existence of an anthraquinone molecule possessing a carboxyl group, hydroxyl group, a terminal methyl group along with a non-hydroxyl fatty acyl chain (Eyong et al., 2005; Luan et al., 2016). Structural characterization of the OKS–PKR reaction product further clarified its chemical identity as 2-carboxy anthraquinone. As per the previous report by (Abe et al., 2005a), heterologously expressed (*E. coli*) OKS from *Aloe arborescens* produced SEK4/SEK4b. Moreover, (Abe et al., 2005b) hypothesized the formation of chrysophanol anthrone (1,8-dihydroxy-3-methyl-9-oxoanthracene-2-carboxylic acid or C_15_H_12_0_3_) through the combined action of OKS and PKR. The theory underlying the proposed approach appears to be logical, considering the suggested structure. However, their nomenclature is not precise or consistent with accepted scientific terminology. According to the National Library of Medicine (NLM, NCBI), chrysophanol anthrone implies 1,8-dihydroxy-3-methyl-10H-anthracen-9-one, (https://pubchem.ncbi.nlm.nih.gov/compound/Chrysophanol-9-anthrone). Our findings helps in distinguishing the product from compounds previously described as chrysophanol anthrone in the literature and pioneered the formation of 2-carboxyl anthraquinone as a cumulative product of OKS and PKR under standard conditions.

The present study also validated the cytoplasmic localization of the *OKS* gene in *Nicotiana benthamiana* (tobacco) leaves through transient expression. The transient approach has been proven effective in studies related to gene function assignment (Johansen and Carrington, 2001; Wroblewski et al., 2005), promoter element analysis (Hellens et al., 2005), and inducible gene expression (Lee and Yang, 2006). Additionally, considering the limited understanding of *OKS* and *PKR* genes in the context of aloin biosynthesis and the robustness of the CRISPR/Cas9 system for gene editing, it has been proven to be an innovative and valuable strategy for investigating the roles of *OKS* and *PKR* in this biosynthetic process. These findings indicated that the *OKS* and *PKR* genes have a crucial role in aloin biosynthesis in *Aloe vera*. Targeted genome editing is a modern technique used to alter the genetic makeup of organisms, and its implementation has been documented in diverse species. Genome editing can be employed to gain insights into gene function or create novel characteristics to enhance crop quality (Kaur et al., 2018). Conclusively, our results suggest that the *OKS* and *PKR* genes play a significant role in aloin/anthraquinone biosynthesis, and these genes could be potential targets for engineering the *Aloe vera* genome to regulate the aloin level and enhance the pharmacological value of aloe plants. The present study explores the pivotal role of OKS and PKR genes in the anthraquinone biosynthesis pathway of *Aloe vera*, employing both heterologous gene expression in *E. coli* and CRISPR/Cas9-mediated genome editing in *Aloe vera*.

## Supplementary data

The following supplementary data are available at JXB online.

Table S1: List of various monocot and dicot plant species selected for phylogenetic analysis of OKS protein

Table S2: List of gene-specific primers used for heterologous expression of OKS gene in *E. coli*

Table S3: Oligonucleotides used for the preparation of oligo-duplex

Table S4: The primer sequence used for screening of transformants

Table S5: Primer sequence used for subcellular localization of OKS gene in the leaves of *Nicotiana benthamiana* L. (tobacco)

Fig. S1: Conserved domain analysis of OKS protein showing the presence of conserved catalytic tetrad of the chalcone synthase superfamily

Fig. S2: Alignment of aloe-based OKS protein sequence with previously reported OKS/PKS protein sequences of *Aloe arborescens*, *Zea mays, Arabidopsis thaliana, Hordeum vulgare, Avena sativa,* and *Sorghum bicolor.* The highly conserved regions are marked in different colours, while the conserved catalytic regions are marked in the black box

## Acknowledgements

The authors would like to thank Chaudhary Charan Singh Haryana Agricultural University, Hisar (Haryana), Guru Jambheshwar University of Science and Technology, Hisar (Haryana) and BRIC-National Agri-Food and Biomanufacturing Institute, Mohali (Punjab), India, for providing the necessary facilities and a suitable environment for executing the present study.

## Author contributions

VC and AJ conceptualized the article and designed the experiments. AJ conducted experiments and prepared the original draft. AJ and ST contributed to data interpretation and curation. VC contributed to the proofreading of manuscript. All authors have read and approved the final manuscript.

## Data availability

The authors declare that all data supporting the findings of this study are available within the article. The nucleotide and deduced amino acid sequence of *OKS* and *PKR* genes have been submitted to the NCBI database (https://www.ncbi.nlm.nih.gov/) via GenBank accession number BankIt2714563 Locus_3851_ Transcript_1/9 OR146495 and BankIt2620826 AvAKR, respectively. The original transcriptome data set could be accessed through Sequence Read Archive (NCBI) via accession number SRR5167034 (https://www.ncbi.nlm.nih.gov/sra).

